# Plasma plasmalogen levels and risk of lymph node positive breast cancer

**DOI:** 10.64898/2025.12.11.693510

**Authors:** Mahsa Yavari, Kristen D. Brantley, Alanis Carmona, Marie Sabatier, Yanshan Liang, June Monge-Lorenzo, Jordan Torpey, Krystina J. Szylo, Midori Flores, Mario Palma, Cameron S. Fraser, Milena Chaufan, Mayher Kaur, Lewis Hendrianto, Whitney Henry, Sheng Tony Hui, Raphaël Rodriguez, Walter C. Willett, Julian Avila-Pacheco, Clary B. Clish, A. Heather Eliassen, Oana A. Zeleznik, Jessalyn Ubellacker

**Author notes:** Authors contributed equally (Mahsa Yavari, Kristen D. Brantley). Authors contributed equally (Oana A. Zeleznik, Jessalyn Ubellacker).

## Abstract

Lymph node involvement is a key predictor of poor breast cancer prognosis. Systemic lipid alterations can contribute to cancer cell survival in lymph nodes, but their relevance in humans remains unclear. Here, we combine human epidemiologic analyses from the Nurses’ Health Study 2 and complementary mechanistic studies in mouse models to investigate how systemic lipid profiles relate to lymph node positive breast cancer. We show that in pre-diagnostic human plasma (n=511), lower levels of phosphatidylethanolamine (PE) and phosphatidylcholine (PC)-enriched plasmalogens are associated with increased risk of lymph node positivity, with stronger associations in samples collected closer to diagnosis. Consistent with the human data, lower levels of PE and PC-enriched plasma plasmalogens are found in mice with nodal involvement. Furthermore, in mice, dietary PUFA depletion reduces lipid oxidation in breast cancer cells in lymph nodes and promotes their survival and metastatic spread. These findings suggest that reduced levels of PE and PC-plasmalogens and decreased PUFA availability creates a lipid environment that enables breast cancer lymph node involvement.

## INTRODUCTION

Each year, over 40,000 people die from breast cancer in the U.S., with metastasis being the leading cause of mortality^1–3^. Breast cancer cells frequently spread first to regional lymph nodes, making nodal involvement a critical predictor of distant metastasis and death^4–9^. Emerging evidence has shown that adaptations in lipid metabolism can increase survivability of cancers in lymph nodes, such as by upregulation of fatty acid uptake, bile acid synthesis, increased lipid storage, and increased beta oxidation^10–13^. However, we lack a detailed understanding of how systemic lipids affect breast cancer cell survival and lymph node positivity in humans.

Oleic acid, a monounsaturated fatty acid (MUFA) abundant in lymph, can protect cancer cells from lipid oxidation and subsequent cell death^14^, whereas polyunsaturated fatty acids (PUFAs) can promote lipid oxidation and cell death in cancer cells^15^. With regard to protecting cancer cells from lipid oxidation, plasmalogens are a subclass of ether phospholipids that make up a significant portion of cell membranes and are also known to influence membrane structure, redox balance, and immune signaling. Because of their emerging roles in cancer cell vulnerability to oxidative stress and inflammation, we focused on plasmalogens in this study, hypothesizing that their depletion could shape a lipid environment that promotes lymph node involvement.

To advance this understanding, we conducted extensive analyses on plasma lipid profiles measured in a breast cancer case-control study nested within the Nurses’ Health Study 2 (NHS2) to examine their association with lymph node positive breast cancer. The NHS2 is a large, ongoing prospective cohort study that collects detailed lifestyle, dietary, medical information, as well as plasma samples from thousands of women to identify risk factors for chronic diseases, including breast cancer^15–17^. In parallel, we performed complementary mechanistic studies in mouse models. To mirror the variations of systemic lipid content present in the heterogeneous NHS2 study population, we placed cohorts of mice on diets with varying MUFA/PUFA ratios. Finally, we investigated the impact of different plasma and lymph fluid lipid levels on LN positive (LN+) versus LN negative (LN-) breast cancer using mouse models.

## RESULTS

### Characteristics at blood draw and diagnosis among breast cancer cases in NHS2 by lymph node status

The main analytic sample included 511 women in NHS2 diagnosed with stage I-III breast cancer with available prediagnostic plasma metabolomic profiling and fatty acid measurements. A total of 152 cases presented with LN+ breast cancer (**Table 1**). Among LN+ cases, most (74%) had breast cancer with 1 to 3 nodes involved and were diagnosed with Stage 2 disease (67%). Subtype was similarly distributed between LN+ and LN-groups, with HR+/HER2-as the most common subtype (59% LN+, 57% LN-). Age and menopausal status at blood draw was similar across LN+ and LN-groups. The mean age among all cases was 45 years at blood draw (SD=4.4 years), and 86% of cases (n=442) were identified as premenopausal, or of unknown menopausal status (from here on considered as premenopausal), at blood draw. Additionally, 78% of women who developed breast cancer were parous, with the most common group having their first birth at age ≥25 years and having 1-2 children. Key risk factors, such as family history, history of benign breast disease, alcohol intake, BMI at age 18y, and BMI at blood draw, were similar between LN+ and LN-cases.

**Table 1.** Characteristics of NHS2 breast cancer cases with plasma fatty acid and metabolite measurements, by lymph node involvement at time of blood draw (N=511).

|  | Lymph Node Positive<br>(N=152) | Lymph Node Negative<br>(N=359) | Overall<br>(N=511) |
| --- | --- | --- | --- |
| Number of lymph nodes |  |  |  |
| None | 0 (0%) | 359 (100%) | 359 (70.3%) |
| 1-3 | 113 (74.3%) | 0 (0%) | 113 (22.1%) |
| 4-9 | 32 (21.1%) | 0 (0%) | 32 (6.3%) |
| 10+ | 7 (4.6%) | 0 (0%) | 7 (1.4%) |
| Stage at BC diagnosis |  |  |  |
| Stage I | 0 (0%) | 291 (81.1%) | 291 (56.9%) |
| Stage II | 102 (67.1%) | 54 (15.0%) | 156 (30.5%) |
| Stage III | 42 (27.6%) | 0 (0%) | 42 (8.2%) |
| Missing | 8 (5.3%) | 14 (3.9%) | 22 (4.3%) |
| Tumor subtype |  |  |  |
| HR+/HER2- | 90 (59.2%) | 204 (56.8%) | 294 (57.5%) |
| HR+/HER2+ | 20 (13.1%) | 54 (15.0%) | 74 (14.5%) |
| HR-/HER2+ | 7 (4.6%) | 18 (5.0%) | 25 (4.9%) |
| HR-/HER2- | 19 (12.5%) | 39 (10.9%) | 58 (11.4%) |
| Unknown/Missing | 16 (10.5%) | 44 (12.3%) | 60 (11.7%) |
| Age at blood draw (years), mean (SD) | 44.9 (4.42) | 45.4 (4.41) | 45.2 (4.41) |
| Age at BC diagnosis (years), mean (SD) | 50.8 (4.81) | 51.5 (4.99) | 51.3 (4.94) |
| Years from blood draw to diagnosis, mean (SD) | 5.93 (3.51) | 6.10 (3.36) | 6.05 (3.40) |
| Year of blood draw, mean (SD) | 1998 (0.82) | 1998 (0.82) | 1998 (0.82) |
| Time of day of blood draw |  |  |  |
| Midnight-7am | 42 (27.6%) | 85 (23.7%) | 127 (24.9%) |
| 8am-noon | 87 (57.2%) | 239 (66.6%) | 326 (63.8%) |
| 1pm-11pm | 16 (10.5%) | 28 (7.8%) | 44 (8.6%) |
| Missing | 7 (4.6%) | 7 (1.9%) | 14 (2.7%) |
| Fasting status |  |  |  |
| Fasting | 102 (67.1%) | 249 (69.4%) | 351 (68.7%) |
| Non-fasting | 50 (32.9%) | 110 (30.6%) | 160 (31.3%) |
| Menopausal status and PMH use at blood draw |  |  |  |
| Premenopausal | 122 (80.3%) | 258 (71.9%) | 380 (74.4%) |
| Postmenopausal, No PMH | 2 (1.3%) | 6 (1.7%) | 8 (1.6%) |
| Postmenopausal, with PMH use | 13 (8.6%) | 48 (13.4%) | 61 (11.9%) |
| Dubious | 15 (9.9%) | 47 (13.1%) | 62 (12.1%) |
| Missing | 0 (0%) | 0 (0%) | 0 (0%) |
| Race |  |  |  |
| White | 151 (99.3%) | 347 (96.7%) | 498 (97.5%) |
| Black | 0 (0%) | 3 (0.8%) | 3 (0.6%) |
| Asian | 0 (0%) | 6 (1.7%) | 6 (1.2%) |
| Other/missing | 1 (0.7%) | 3 (0.8%) | 4 (0.8%) |
| Ethnicity |  |  |  |
| Hispanic | 3 (2.0%) | 5 (1.4%) | 8 (1.6%) |
| Non-Hispanic | 149 (98.0%) | 354 (98.6%) | 503 (98.4%) |
| Age at menarche (years), mean (SD) | 12.4 (1.22) | 12.3 (1.37) | 12.4 (1.33) |
| Age at first birth and parity |  |  |  |
| Nulliparous | 29 (19.1%) | 82 (22.8%) | 111 (21.7%) |
| 1-2kids <25yr | 18 (11.8%) | 53 (14.8%) | 71 (13.9%) |
| 1-2kids ≥25yr | 66 (43.4%) | 130 (36.2%) | 196 (38.4%) |
| 3+kids <25 | 18 (11.8%) | 44 (12.3%) | 62 (12.1%) |
| 3+kids 25+yr | 21 (13.8%) | 50 (13.9%) | 71 (13.9%) |
| Ever breastfed |  |  |  |
| Yes | 103 (67.8%) | 226 (63.0%) | 329 (64.4%) |
| No | 49 (32.3%) | 133 (37.1%) | 182 (35.6%) |
| Oral contraceptive use |  |  |  |
| Current user | 5 (3.3%) | 13 (3.6%) | 18 (3.5%) |
| Past user | 129 (84.9%) | 291 (81.1%) | 420 (82.2%) |
| Never user | 18 (11.8%) | 55 (15.3%) | 73 (14.3%) |
| Missing | 0 (0%) | 0 (0%) | 0 (0%) |
| Family history of BC |  |  |  |
| Yes | 23 (15.1%) | 55 (15.3%) | 78 (15.3%) |
| No | 129 (84.9%) | 304 (84.7%) | 433 (84.7%) |
| History of benign breast disease |  |  |  |
| Yes | 27 (17.8%) | 85 (23.7%) | 112 (21.9%) |
| No | 125 (82.2%) | 274 (76.3%) | 399 (78.1%) |
| BMI at age 18 (kg/m<sup>2</sup>), mean (SD)* | 20.6 (2.64) | 20.8 (2.66) | 20.7 (2.65) |
| BMI (kg/m<sup>2</sup>) at blood draw, mean (SD) | 25.3 (4.63) | 25.3 (5.44) | 25.3 (5.21) |
| Weight change since age 18 (kg), mean (SD)* | 13.0 (11.0) | 12.3 (12.6) | 12.5 (12.1) |
| Alcohol intake (drinks/week), mean (SD)* | 4.09 (9.62) | 3.57 (5.86) | 3.72 (7.19) |
| Physical activity (met-hours/week), mean (SD)* | 16.4 (15.0) | 18.2 (14.4) | 17.6 (14.6) |
| Total plasma MUFA (% of total FA) | 16.2 (1.40) | 16.3 (1.35) | 16.3 (1.36) |
| Total plasma PUFA (% of total FA) | 37.5 (2.75) | 37.2 (2.83) | 37.3 (2.81) |
| Plasma MUFA:PUFA ratio | 0.437 (0.06) | 0.441 (0.06) | 0.440 (0.06) |

MUFA= monounsaturated fatty acid, PUFA=polyunsaturated fatty acid
\*Missing values imputed with mean value

### Prediagnostic plasma PE and PC plasmalogen levels were lower in women with subsequent LN+ v. LN-breast cancer

While no plasma metabolites appeared to be statistically significantly associated with LN-positivity after adjustment for the number of effective tests (n=215, adjusted p-value for significance=0.0002), we observed strong inverse associations, which were nominally significant, for many phosphatidylethanolamine (PE) and phosphatidylcholine (PC) plasmalogens. Among the 20 individual metabolites that were found to be most strongly associated with LN-positivity, 7 were plasmalogens (**Table 2**). Increases in each of these 7 plasmalogens (from the 10^th^ to 90^th^ percentile), were associated with 50-60% reduced odds of LN+ versus LN-breast cancer. Beyond these top metabolites, plasmalogens tended to be inversely associated with LN+ breast cancer overall (**Table S1**). Individual metabolite results were similar when further adjusting for tumor subtype. We also evaluated the association between an individuals’ average level of PE and PC plasmalogens (calculated by taking the average value after summing of all PE and PCs) with LN+ versus LN-breast cancer. Higher average PE and PC plasmalogen level (10^th^ to 90^th^ percentile contrast) was associated with a statistically significant reduction in odds of LN+ breast cancer of 52% (p=0.03) in the model adjusted for blood-draw factors (**Table 3**). This reduction was similar in magnitude and borderline significant (p=0.05) in the fully adjusted model (**Table 3**). Inverse associations were suggestively stronger among those with blood draw closer to time of diagnosis for both individual (**Table S2**) and average (**Table 3)** plasmalogen levels, though the interaction term was not statistically significant (p=0.65). In addition, we found the inverse association between average plasma plasmalogens and LN-positivity was suggestively stronger for those individuals with BMI <25 kg/m2 [adjusted OR (95% CI) BMI <25 kg/m^2^=0.43 (0.16-1.12) vs. BMI ≥25 kg/m^2^= 0.60 (0.19-1.83), p-interaction=0.62] (**Table S3**).

**Table 2.**
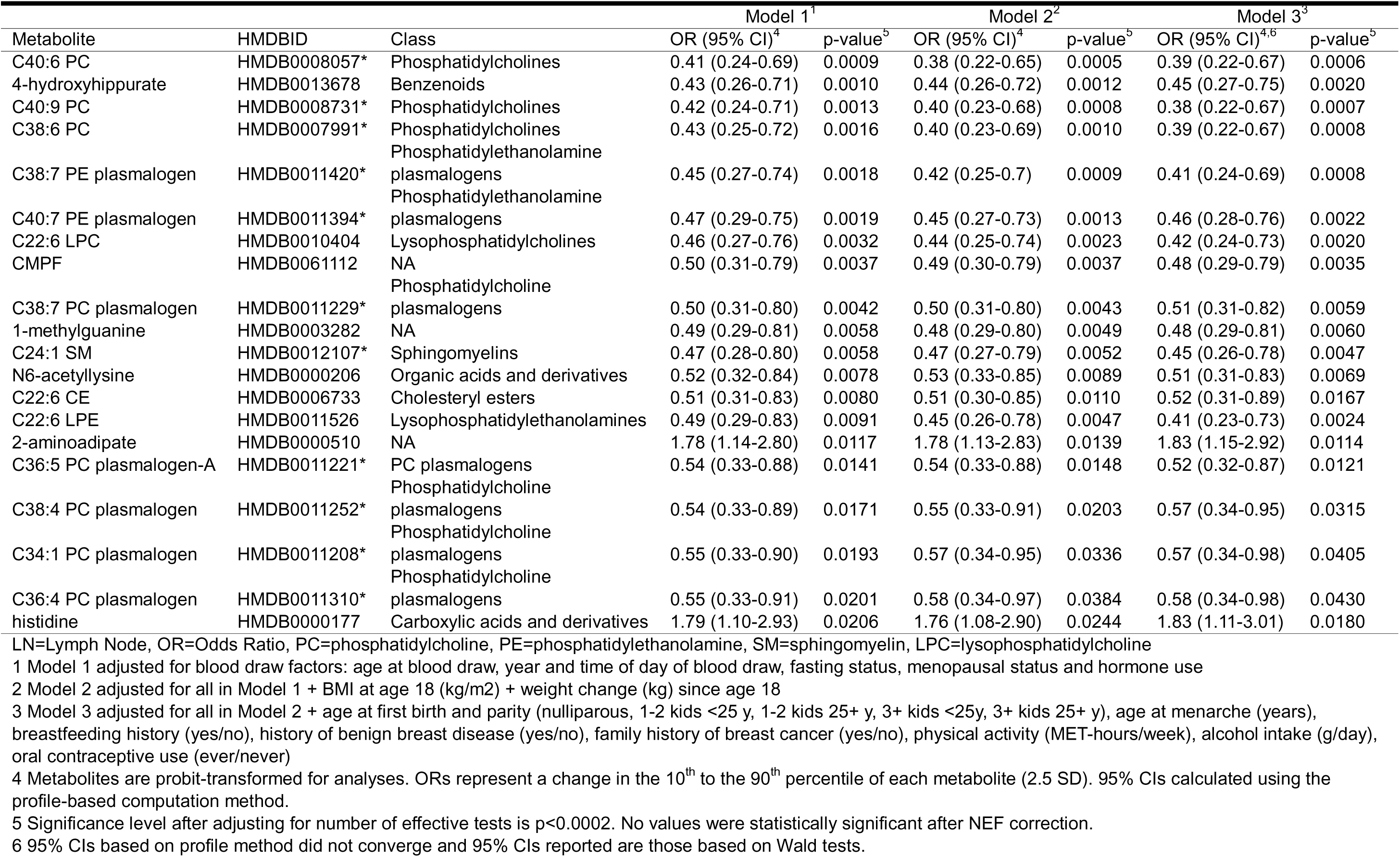
Top 20 Metabolite Associations with LN Positive (vs. LN Negative) Breast Cancer in NHS2 Subset (N=511).

**Table 3.** Association between average plasmalogen level and LN positive vs. LN negative BC, overall and by time from blood draw to diagnosis.

| Subset | LN+/LN- | Model 1<sup>1</sup> |  | Model 3<sup>2</sup> |  |
| --- | --- | --- | --- | --- | --- |
|  |  | OR (95% CI)<sup>3</sup> | p-value | OR (95% CI)<sup>3</sup> | p-value |
| Overall | 152/359 | 0.48 (0.24-0.94) | 0.03 | 0.50 (0.25-1.00) | 0.05 |
| Years, blood draw to dx |  |  |  |  |  |
| <6.25 y | 74/181 | 0.37 (0.14-0.97) | 0.05 | 0.49 (0.17-1.36) | 0.18 |
| ≥6.25 y | 78/178 | 0.64 (0.24-1.67) | 0.36 | 0.52 (0.18-1.44) | 0.21 |

LN=lymph node, OR=odds ratio
All metabolites were probit-transformed and average PE plasmalogen + PC plasmalogen was taken for each individual.
1 Model 1 is adjusted for all blood draw factors: age at blood draw, year and time of day of blood draw (midnight-7am, 8am-noon, 1pm-11pm), fasting status, menopausal status and hormone use
2 Model 3 is adjusted for all blood draw factors + BMI at age 18 (kg/m<sup>2</sup>) + weight change (kg) since age 18 + age at first birth and parity (nulliparous, 1-2 kids <25 y, 1-2 kids 25+ y, 3+ kids <25y, 3+ kids 25+ y), age at menarche (years), breastfeeding history (yes/no), history of benign breast disease (yes/no), family history of breast cancer (yes/no), physical activity (MET-hours/week), alcohol intake (g/day), oral contraceptive use (ever/never)
3 ORs represent an increase in average plasma plasmalogen level from the 10<sup>th</sup> to 90<sup>th</sup> percentile (2.5 SD)

Findings from individual and average PC and PE plasmalogens were further supported by results from a metabolite set enrichment analysis. When grouping metabolites by structural classes (N=19), we found that PC plasmalogens and PE plasmalogens were negatively enriched among LN+ breast cancer cases relative to other metabolite classes (**Figure 1**), which was consistent across different adjustment models (**Figure S1, Table S4**). PC plasmalogens had a normalized enrichment score (NES) of −1.68 (p_adj_=0.04) and PE plasmalogens had a NES of −1.41 (p_adj_=0.24) in fully adjusted models (**Table S4**). Associations were strongest for those with blood draws closest to diagnosis (**Figure 1, Figure S2**, **Table S5)**. For example, comparing those with <6.25 years to those with >6.25 years between blood draw to diagnosis, for PC plasmalogens NES= −1.87 ( (p_adj_=0.01) vs. −1.43 (pad=0.13) and for PE plasmalogens NES=-1.89 (p_adj_=0.01) vs. −1.44 (p_adj_=0.12) (Table S5). Plasmalogen metabolite group associations were also strongest among individuals with BMI <25 kg/m^2^ (LN+ N=82, LN-N=209) compared to those with BMI ≥25 kg/m^2^ (LN+ N=70, LN-N=150) (e.g., PC plasmalogens NES= −1.92 (p_adj_=0.01) vs. −1.28 (p_adj_=0.33)(Table S5, **Figure S3**).

**Figure 1.**
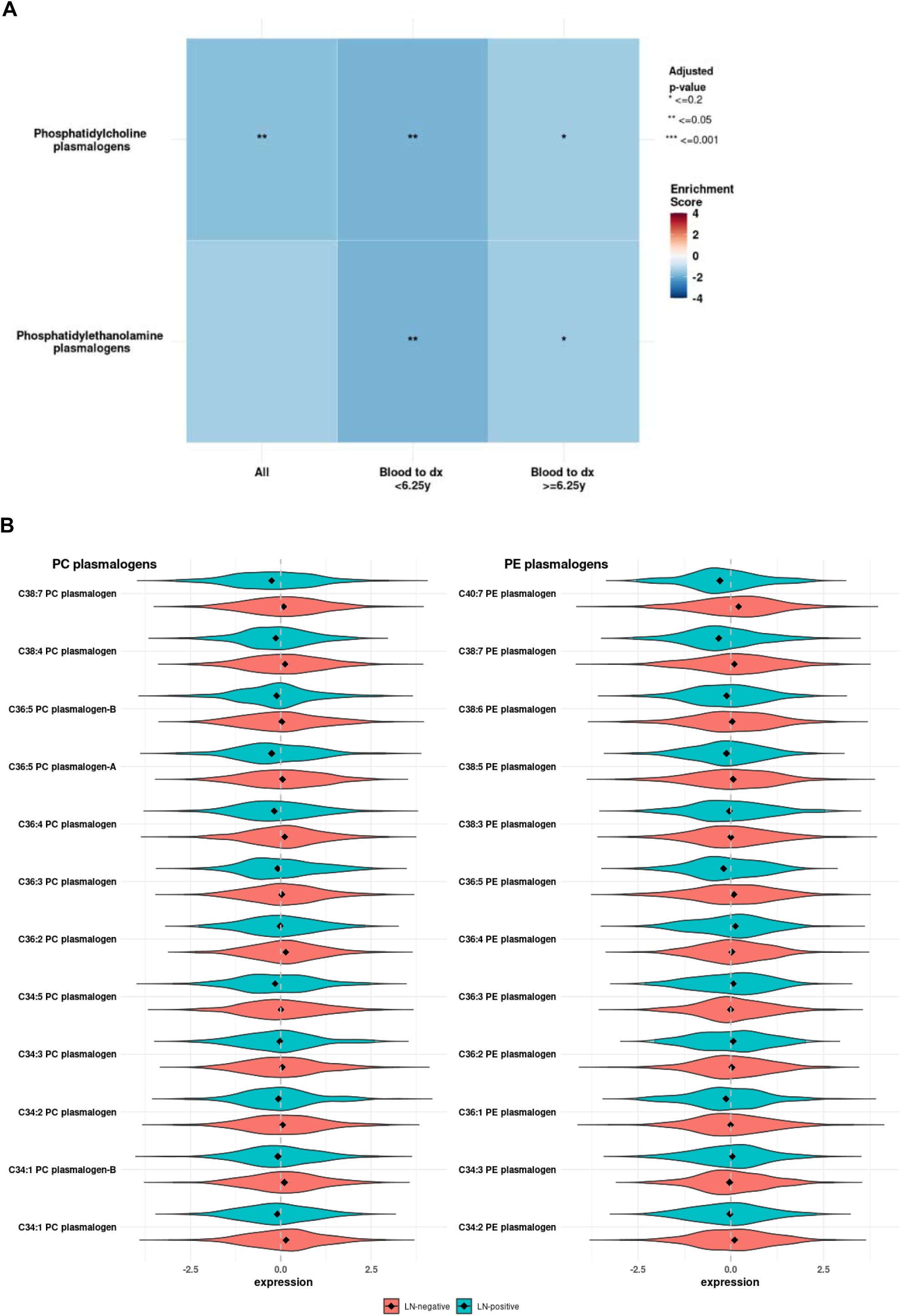
Relative enrichment and expression of plasmalogen metabolites and pathways in lymph node-positive (LN+) compared to lymph node-negative (LN-) breast cancer overall. (N=511, LN+=152, LN-=359). **(A)** Relative enrichment in plasmalogen metabolite pathways by years from blood draw to diagnosis (below or above median time). Metabolite subclasses were defined by the Broad Institute based on structural similarities. Models were adjusted for blood draw factors (age at blood draw, year of blood draw, time of day of blood draw, fasting status at blood draw, menopausal status and postmenopausal hormone use at blood draw), BMI at age 18 (kg/m2), weight change since age 18 (kg), age at menarche (years), age at first birth and parity (nulliparous, <25y and 1-2 births, <25y and 3+ births, ≥25y and 1-2 births, ≥25y and 3+ births), family history of breast cancer, history of benign breast disease, average alcohol intake (g/day), physical activity (met-hours/week), breastfeeding (ever v. never), oral contraceptive use (ever v. never). **(B)** Distribution of probit-transformed plasmalogen expression within each subclass (PC or PE plasmalogen) by LN status (turquoise=LN-positive, pink=LN-negative). Median expression values for each plasmalogen are shown by the black diamond inside violin plots.

### Plasma MUFA/PUFA ratios did not differ between LN+ and LN-breast cancer cases

Plasma MUFA/PUFA ratios were not associated with LN+ vs. LN-breast cancer overall (**Table 4**). The non-significant relationship was maintained when examining associations based on timing between blood draw and diagnosis (Table 4, Table S6) and based on BMI at blood draw (Table S6).

**Table 4.** Association between plasma MUFA:PUFA ratio and LN positive vs. LN negative breast cancer, overall and by time from blood draw to diagnosis.

| Subset | LN+/LN- | Model 1<sup>1</sup> |  | Model 3<sup>2</sup> |  |
| --- | --- | --- | --- | --- | --- |
|  |  | OR (95% CI)<sup>3</sup> | p-value | OR (95% CI)<sup>3</sup> | p-value |
| Overall | 152/359 | 0.89 (0.56-1.40) | 0.63 | 0.84 (0.52-1.34) | 0.48 |
| Years, blood draw to dx |  |  |  |  |  |
| <6.25 y | 74/181 | 1.10 (0.58-2.04) | 0.76 | 1.03 (0.52-1.95) | 0.93 |
| ≥6.25 y | 78/178 | 0.74 (0.34-1.56) | 0.44 | 0.68 (0.29-1.53) | 0.36 |

LN=lymph node, OR=odds ratio
1 Model 1 is adjusted for all blood draw factors: age at blood draw, year and time of day of blood draw (midnight-7am, 8am-noon, 1pm-11pm), fasting status, menopausal status and hormone use
2 Model 3 is adjusted for all blood draw factors + BMI at age 18 (kg/m<sup>2</sup>) + weight change (kg) since age 18 + age at first birth and parity (nulliparous, 1-2 kids <25 y, 1-2 kids 25+ y, 3+ kids <25y, 3+ kids 25+ y), age at menarche (years), breastfeeding history (yes/no), history of benign breast disease (yes/no), family history of breast cancer (yes/no), physical activity (MET-hours/week), alcohol intake (g/day), oral contraceptive use (ever/never)
3 ORs represent an increase in MUFA:PUFA from the 10<sup>th</sup> to 90<sup>th</sup> percentile ( $\Delta$ +0.137)

### Restriction to premenopausal individuals at blood draw and diagnosis

Though the majority (86%) of individuals in NHS2 were premenopausal at blood draw, we conducted sensitivity analyses to determine the influence of postmenopausal status on our results. When restricting to women premenopausal at blood draw only (n=442), our results were comparable to those presented among all women for both plasmalogen (Table S7) and MUFA:PUFA (Table S8) associations with LN+ breast cancer. Further restriction to individuals premenopausal at both blood draw and diagnosis (N=242) produced similar results, though resulting subsets were small, leading to larger confidence intervals (**Table S7, Table S8**).

To summarize the key results from the NHS2 cohort studies, C38:7 PE plasmalogen, and C40:7 PE plasmalogen were nominally significantly lower in prediagnostic plasma samples from participants who later developed LN+ compared to LN-breast cancer. When grouping metabolite classes, PE and PC plasmalogens were strongly inversely associated with LN-positivity (**Figure 1**). Plasma MUFA/PUFA ratios were not statistically significantly associated with LN-positivity, though different trends in associations were apparent based on time from blood draw to diagnosis and BMI at blood draw.

Next, to test the biological relevance of our epidemiologic findings and explore causal mechanisms, we conducted controlled dietary studies in mouse models, using defined MUFA/PUFA diets to mimic the variation in systemic lipid profiles observed in the plasma samples from the NHS2 population.

### Mouse models of LN+ and LN-breast cancer with varied plasma lipid profiles

We designed four diets with varying MUFA/PUFA ratios by adjusting the proportions of specific fatty acids in each diet. The MUFA/PUFA composition of each diet was controlled by incorporating oils with distinct fatty acid profiles (**Table S9**). Confounding factors were minimized by isocalorically matching all diets and maintaining total fat content at 25% of calories. We refer the four diets as Q1-4, based on their MUFA/PUFA ratios per gram of oil: Q1 (0.1), Q2 (0.4), Q3 (3.3), and Q4 (5.5) (**Fig. 2a**).

**Figure 2.**
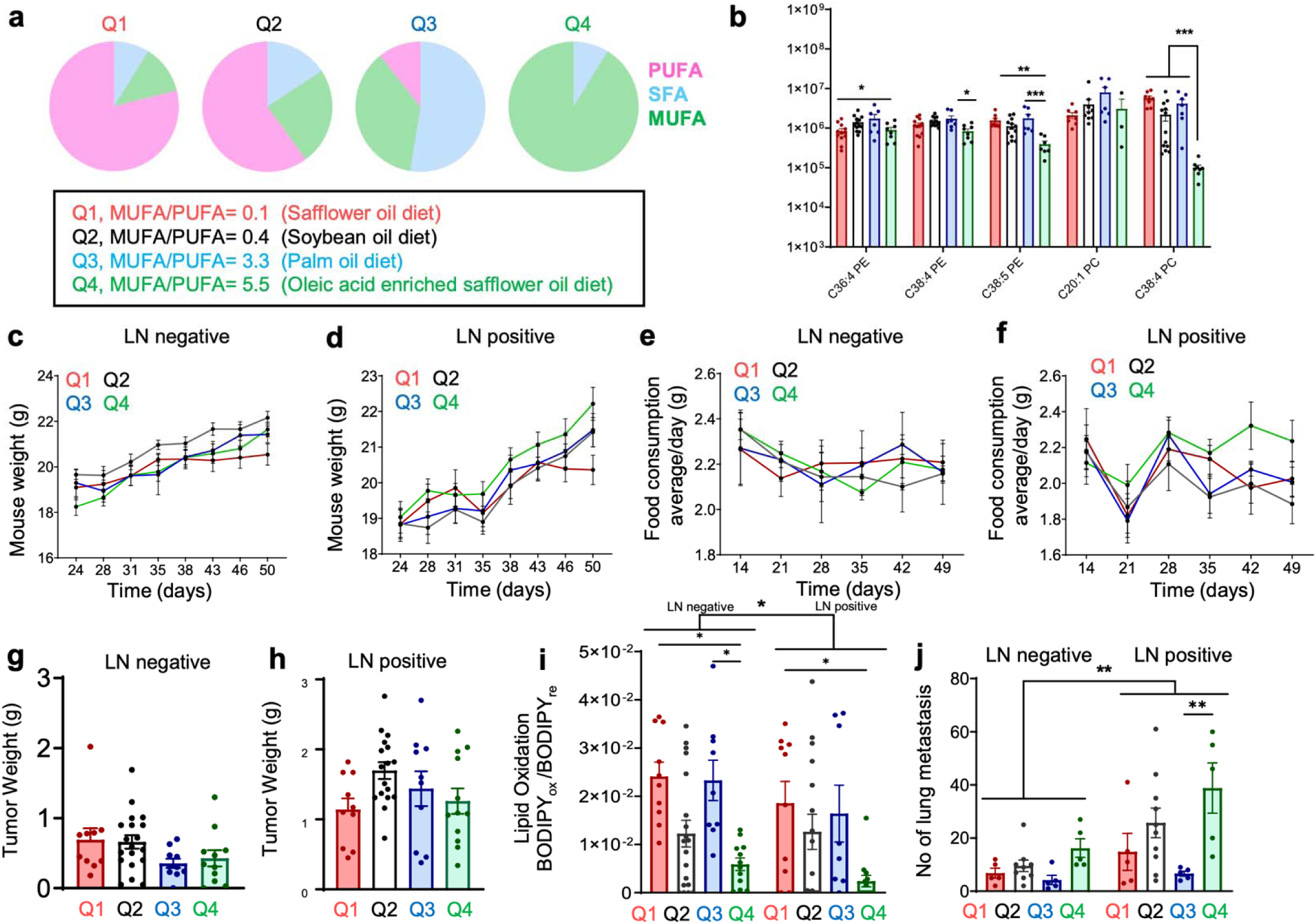
Mice with increased MUFA/PUFA intake and bearing 4T1 breast cancer in the mammary fat pad (LN negative) or popliteal lymph node (LN positive) have decreased plasma plasmalogens, increased metastasis, and decreased lipid peroxidation levels in cancer cells. (**a**) Composition of fatty acids in 4 generated diets (Q1, Q2, Q3, Q4) formulated to mimic differing MUFA/PUFA intake levels. (**b**) Plasmalogen PE and PC differences in plasma of different diet groups. (**c,d**), food consumption (**e,f**), were monitored. At the experimental endpoint, final tumor weights were obtained (**g,h**). (**i**) At the experimental endpoint, tumor cells from the mammary fat pad (LN negative) or popliteal lymph nodes (LN positive) cohorts were isolated, and lipid peroxidation was measured by flow cytometry analysis of BODIPY-C11 staining. (**j**) Lung metastasis forming after mammary fat pad (**j**) or from the popliteal lymph node injection were detected by H&E quantifications of the presence of metastasis (see methods for details on quantification). For all panels, each bar represents the mean ± SD. Data from at least 2 independent in vivo experiments were analyzed by one-way ANOVA or 2-way ANOVA in LN positive and LN negative combined graphs followed by Tukey’s multiple comparison.*p <0.05, **p <0.01, ***p <0.001.

To model breast cancer in the mouse studies, we used the triple-negative 4T1 mouse cell line. This line spontaneously metastasizes after primary injection into the mammary fat pad, thus mimicking breast cancer development and progression in humans. Syngeneic Balb/c mice were fed the Q1-Q4 diets for six weeks. After two weeks on the diet, half of the mice in each diet group were injected with 4T1 cells either orthotopically (into the mammary fat pad) or intranodally (into the popliteal lymph node) to model differences in LN positivity. Orthotopic injections represent LN-breast cancer, as mice in these groups had tumor formation at the primary site but not the lymph nodes, whereas intranodal injections represent LN+ disease. Once the tumors reached approximately 2.0 cm in diameter, the mice were euthanized.

### Plasma PE(38:4) plasmalogen levels were lower in mice consuming diets with higher MUFA/PUFA ratios

At the experimental endpoint, to assess whether these diets captured the range of plasmalogen differences observed in the NHS2 studies, we collected plasma and lymph fluid from the mice and performed unbiased lipidomic profiling. Levels of C38:4 PE, C20:1 PC, and C38:4 PC were significantly lower in the plasma of the lymph node-positive group compared to the lymph node-negative group (**Figure S4a**). Plasmalogens, specifically PE(38:4), were significantly lower in the plasma of mice on the highest MUFA/PUFA diet (Q4) compared to the other diets groups (**Figure 2b**).These analyses further revealed significant differences in overall lipid composition between plasma and lymph fluid (**Figure S4c**), as well as distinct differences in the lipid profiles of the mice on different diets (**Figure S4d**). Additionally, lipidomic analysis of lymph fluid collected from mouse lymph nodes showed a marked reduction in C36:4 PE, C38:4 PE, and C36:5 PE in the lymph node-positive group (**Figure S4b**). These findings suggest a link between decreased plasmalogen levels and lymph node metastasis, highlighting their possible role in cancer progression and metastatic spread in vivo.

### Mouse models of LN+ and LN-breast cancer with varied plasma lipid profiles had no overall differences in primary tumor growth

No statistically significant differences in mouse weight (**Figure 2c, 2d**), tumor growth (**Figure S4e, S4f**) or food consumption (**Figure 2e, 2f**) were observed across the different mouse diet groups, both for the LN- or LN+ groups. Of note, in the LN+ mouse group, there was a decrease in food consumption on day 21 which can likely be attributed to decreased food consumption typically observed in mice post intranodal injection, which is a minimally invasive surgical procedure (**Figure 2f**). Furthermore, there were no statistically significant differences in the final tumor weights from the different mouse diet groups, both for the LN- or LN+ groups (**Figure 2g, 2h**).

### Breast cancer cells from mice consuming diets with higher MUFA/PUFA ratios had decreased levels of lipid oxidation

PUFAs, unlike MUFAs, contain bis-allylic bonds that renders them more prone to oxidative damage when incorporated into cell membranes^19–21^. Cancer cells have been shown to take up exogeneous lipids available in their microenvironment for incorporation into their cell membranes^14^, and thus we reasoned that breast cancer cells from mice on diets containing more PUFAs might experience more lipid oxidation. To test this, tumor cells from the mammary fat pad (LN-) or popliteal lymph nodes (LN+) groups were isolated, and lipid oxidation was measured by flow cytometry analysis of BODIPY-C11 staining. Overall, the LN+ compared to LN-cohorts had decreased levels of lipid oxidation for each diet cohort, except for the Q2 diet (**Figure 2i**). Tumors isolated from both the mammary fat pads (LN-group) or from the lymph nodes (LN+ group) of mice on the Q4 diet, which had higher levels of MUFA compared to PUFA, had significantly decreased levels of lipid oxidation as measured by BODIPY-C11. This suggests the different MUFA/PUFA dietary intake of the mice leads to alterations in the cancer cells that reduce lipid oxidation levels in mice consuming diets with the highest MUFA/PUFA ratios (Q4).

### Breast cancer cells from mice consuming diets with higher MUFA/PUFA ratios had increased levels of lung metastasis

To determine if these breast cancer cells from mice in the Q4 groups might be better able to survive to form distant metastasis, we next analyzed the lungs from the mouse studies for the presence of spontaneous metastases by pathological quantification of breast cancer cells on H&E tissue sections (**Figure S4g**). No differences in overall lung metastases were observed among mice on the different dietary cohorts in the LN-cohorts, but in the LN+ cohorts, mice consuming the Q4 diet had significantly increased lung metastases compared to Q3 diet (p=0.0017), (**Figure 2j**). Although the mechanism underlying these observed changes is beyond the scope of this current work, these data suggest that the lymph node microenvironment confers lipid-associated alterations on the metastatic potential of breast cancer cells in line with prior studies demonstrating this effect^13,22^.

### In vivo analysis of lipidomic MUFA/PUFA ratios in plasma and breast cancer progression in the 4T1 immunocompetent mouse model of breast cancer

To further explore differences observed in the LN- and LN+ mouse cohorts, we next analyzed all mice from our experimental groups using similar analyses as the NHS2 studies. Plasma from all mice in the dietary cohorts were analyzed for MUFA/PUFA content by LC-MS/MS and mice in the cohort were divided into three tertiles (T1, T2, T3) for analysis based on detected MUFA/PUFA ratios in the plasma to consider the *de novo* effects of lipid metabolism on lipidomic profile of each mouse. When the mice in these groups were divided into these tertiles (instead of by the dietary-based cohorts described in Figure 2), there were again no overall differences in tumor growth or tumor weight at endpoint for either the LN negative cohorts (**Figure 3a, 3b**) or LN positive cohorts (**Figure 3e, 3f**). Tumors isolated from both the mammary fat pads (from LN negative cohorts) or from the lymph nodes (from the LN positive cohorts) of mice in the T3 group, which had higher levels of MUFA compared to PUFA, had significantly decreased levels of lipid oxidation as measured by BODIPY-C11(**Figure 3c, 3g**). This, together with the data in Figure 2i showing the Q4 dietary cohort of mice had tumors with lower lipid oxidation levels compared to other mouse diet groups, further suggests that different MUFA/PUFA dietary intake of the mice impact alterations in the cancer cells resulting in reduced lipid oxidation levels at the highest MUFA/PUFA ratios. No differences in overall lung metastasis were observed among mice on the different tertiles (T1, T2, T3) in the LN negative or LN positive cohorts, although mice in the T3 tertile had a tendency toward an increased number of lung metastasis (**Figure 3d, 3h**).

**Figure 3.**
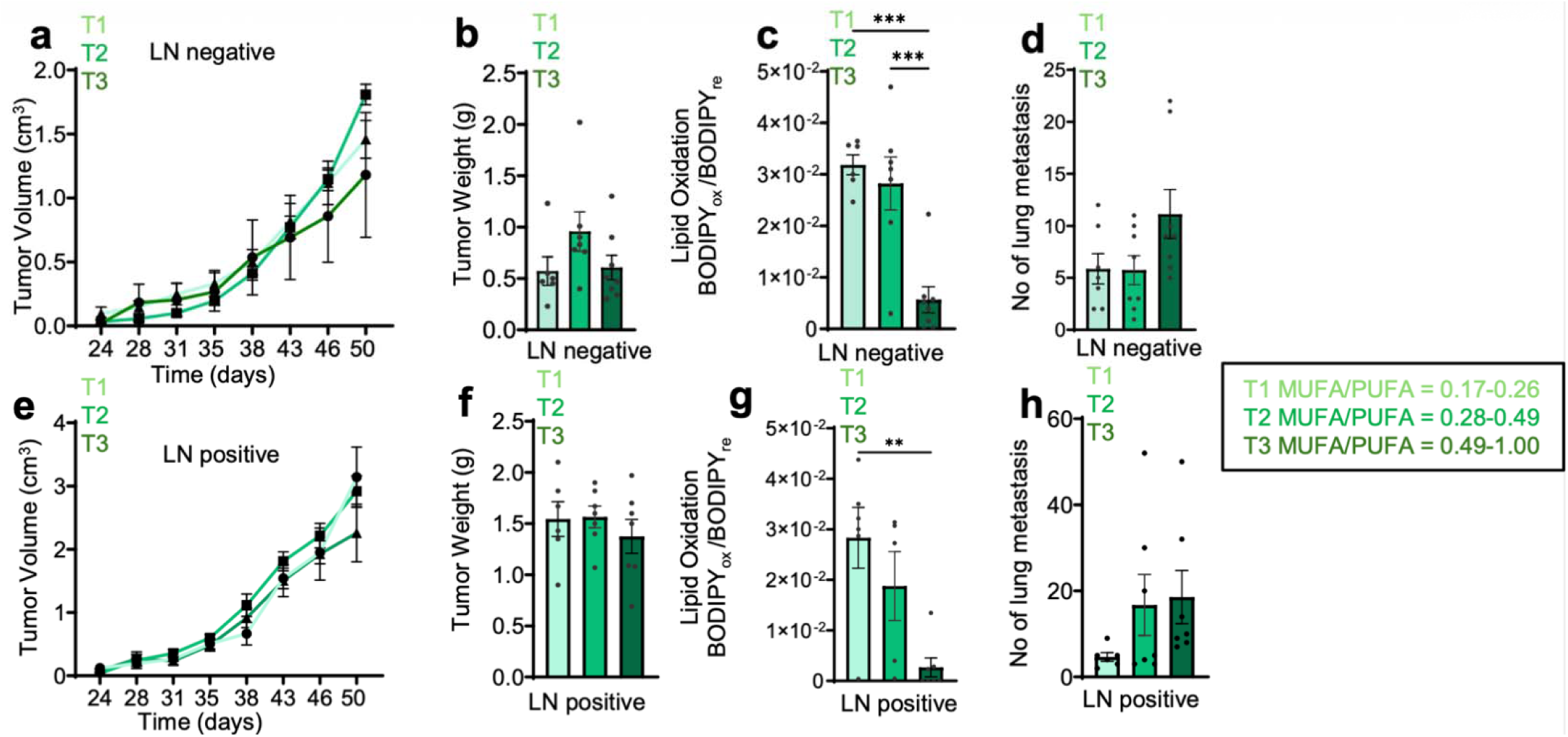
In vivo analysis of lipidomic MUFA/PUFA ratios in plasma progression in the 4T1 immunocompetent mouse model of breast cancer. Plasma from all mice in the dietary cohorts were analyzed for MUFA/PUFA content by LC-MS/MS and mice in the cohort were divided into three tertiles (T1, T2, T3) for analysis based on detected MUFA/PUFA ratios in the plasma, to mirror the analyses performed on the human plasma samples in NHS2. (**a**) tumor growth curves of Balb/c mice injected with 4T1 tumors into the mammary fat pad (LN negative) and (**c**) tumor growth curves of Balb/c mice injected with 4T1 tumors into the popliteal lymph node (LN positive) stratified by MUFA/PUFA tertiles. At the experimental endpoint, final tumor weights were obtained (**b,f**). At the experimental endpoint, tumor cells from the mammary fat pad (LN negative) (**c**) or popliteal lymph nodes (LN positive) (**g**) cohorts were isolated, and lipid peroxidation was measured by flow cytometry analysis of BODIPY-C11 staining. (**d**) Lung metastasis forming after mammary fat pad (**d**) or from the popliteal lymph node (**h**) injection were detected by H&E quantifications of the presence of metastasis (see methods for details on quantification). For all panels, each bar represents the mean ± SD. Data from at least 2 independent in vivo experiments were analyzed by one-way ANOVA or 2-way ANOVA in LN positive and LN negative combined graphs followed by Tukey’s multiple comparison. *p <0.05, **p <0.01, ***p <0.001, ****p <0.0001.

## DISCUSSION

In this study, we combined human epidemiological and mouse model mechanistic data to evaluate the role of lipids in the development of LN+ breast cancer. We evaluated differences in pre-diagnostic lipid profiles between LN+ and LN-breast cancer cases in NHS2 and identified several lipid groups that were associated with LN positivity. We also determined that lymph node status was not associated with plasma MUFA/PUFA ratios assessed on average 6 years before diagnosis. Consistent with the human data, PE and PC plasmalogen levels in plasma and lymph fluid were lower in mice with nodal involvement. Upon finding that PC plasmalogens and PE plasmalogens were inversely associated with LN positivity, we focused on investigating mechanistic connections between lipids and LN positivity in mouse models of metastatic breast cancer. Additionally, after feeding mice with different diets, which reflect the different MUFA/PUFA plasma levels in women from the NHS2, we found breast cancer cells taken from lymph nodes of mice on the lowest MUFA/PUFA diet had increased levels of lipid oxidation, decreased survival in lymph nodes, and decreased formation of lung metastases. These findings suggest that reduced systemic PE and PC-plasmalogens and decreased PUFA availability creates a lipid environment that enables breast cancer lymph node involvement (**Figure 4**). To our knowledge, no prior epidemiologic studies have specifically examined plasmalogens in relation to lymph node status in breast cancer, further emphasizing the novelty of our findings.

**Figure 4.**
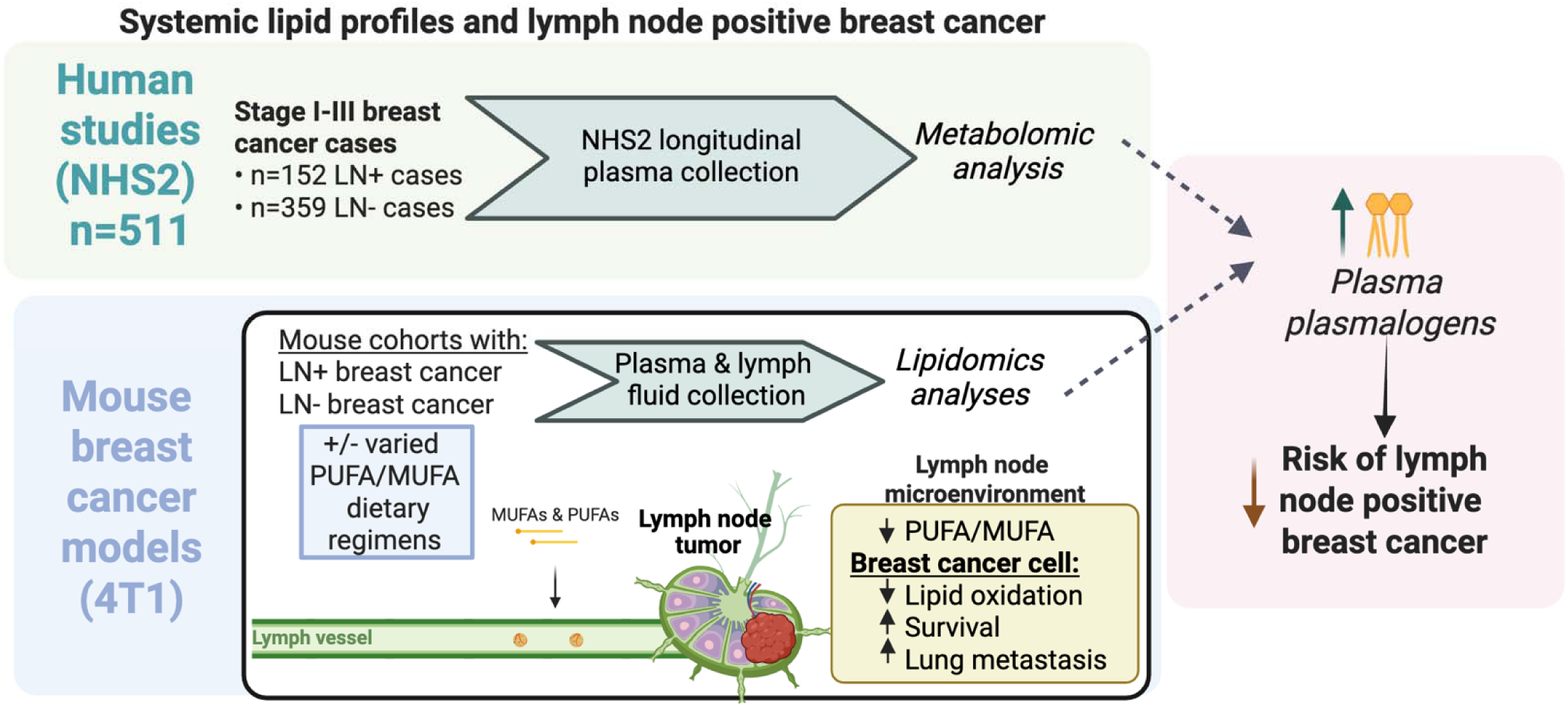
Graphical schematic. Here, we combine human epidemiologic analyses from the Nurses’ Health Study 2 and complementary mechanistic studies in mouse models to investigate how systemic lipid profiles relate to lymph node positive breast cancer. We show that in pre-diagnostic human plasma (n=511), lower levels of PE and PC-enriched plasmalogens are associated with increased risk of lymph node positivity, with stronger associations in samples collected closer to diagnosis. Consistent with the human data, lower levels of PE and PC-enriched plasmalogens are found in mice with nodal involvement. Furthermore, in mice, dietary PUFA depletion reduces lipid oxidation in breast cancer cells in lymph nodes and promotes their survival and metastatic spread. These findings suggest that reduced levels of PE and PC-plasmalogens and decreased PUFA availability creates a lipid environment that enables breast cancer lymph node involvement.

This study has several limitations. Within the NHS2 study, plasma samples were only available for one time-point per participant, limiting our ability to assess the influence of MUFA/PUFA ratios and metabolite levels on LN-positivity over time. It is possible that we are not capturing the time period that is most relevant to the associations explored here. However, we addressed this in part by stratifying analyses by time from blood draw to diagnosis, finding greater associations with lymph node status in periods closer to diagnosis. Given the relatively small numbers of LN+ and LN-BC patients in the case-control subset used here, we were not able to assess the impact of dietary MUFA and PUFA intake levels on LN-positivity, though this is an area for future research. Given that some of our diet formulations, such as Q1 and Q2, included soybean oil rich n-3 PUFAs, it is important to note that while these fatty acids are essential and their influence on tumor progression may be shaped by distinct inflammatory phenotypes, as n-3 or n-6 PUFAs can be precursors to both pro- and anti-inflammatory mediators depending on enzymatic context, tissue environment, and oxidation state.

Our mechanistic mouse models focused on triple-negative breast cancer (TNBC), the subtype most sensitive to lipid oxidation^23^, while most NHS2 cases had estrogen receptor positive breast cancer. This difference in tumor subtype complicates direct comparison between models but allows for complementary strengths—mechanistic insight in TNBC and broader epidemiologic relevance in more common breast cancer subtypes. Additionally, because NHS2 plasma volumes were limited, semi-targeted metabolomic profiling prioritized breadth over molecular resolution. As a result, lipid classes and total double bond content were quantified, but the specific fatty acid chains within each phospholipid or triglyceride were not resolved and thus our interpretations are limited to broader unresolved lipid classes. Finally, it is important to note that NHS2 participants are majority White, which limits the generalizability of our findings to other racial and ethnic populations. Strengths of this study most notably include the use of cohort based epidemiologic data to inform the design of mouse studies to uncover causal mechanisms that could underlie associations between lipids and lymph node positivity in human breast cancer patients. This is particularly advantageous given that lipid studies in human patient samples are often confounded by variables such as patient weight loss, associated fatigue, changes in insulin and glucose signaling, among others.

Plasmalogens, which constitute approximately 20% of total phospholipids in mammals, play a critical role in cell membranes^24^. Elevated plasmalogen levels are present in mammalian cells and tissues such as the brain, heart, testes, and circulating immune cells^25^. Plasmalogens can be synthesized endogenously or exogenously increased via the diet, with particularly high concentrations found in seafood and pork, chicken, or beef^26^. Recent studies further emphasize the significance of plasmalogens in inhibiting systemic inflammation in chronic disease, neuroinflammation, as well as oxidative vulnerability of cancer cells^27–31^. Zou et al showed that plasmalogens lipids are also vulnerable to oxidation in human renal and ovarian carcinoma cells, which triggers death by lipid oxidation (ferroptosis)^29^. They also demonstrate that ether lipids containing polyunsaturated fatty acids (PUFA-ePLs) were essential for this cell death by lipid oxidation. Overall, our findings suggest that reduced systemic levels of PE and PC plasmalogens may create a lipid environment that supports lymph node involvement in breast cancer. Together, our human and mouse data point to a potentially actionable metabolic vulnerability where restoring or preserving plasmalogen levels and PUFA availability could limit metastatic progression in lymph nodes.

## METHODS

### NHS2 Analyses

#### Cohort

The Nurses’ Health Study 2 (NHS2) is an ongoing prospective cohort of 116,429 female nurses aged 25-42 years at enrollment in 1989 and has been described in detail elsewhere^32^. Briefly, participants receive follow-up questionnaires every two years and self-report risk factor information and disease diagnoses. Cases and tumor information are subsequently confirmed by medical record review. Blood samples were collected between 1996-1999 from 29,611 NHS2 participants ages 32-54 years, and collection has been described previously^33^. Briefly, a standard questionnaire was completed at blood collection, indicating the date and time of the collection, hours since the participant last ate, current weight, menopausal status, and medication use. Blood samples were shipped overnight with an icepack and then processed and stored in liquid nitrogen freezers (≤ −130°C). The study protocol was approved by the institutional review boards of the Brigham and Women’s Hospital and Harvard T.H. Chan School of Public Health, and those of participating registries as required.

#### Analytic Sample

Eligible cases included women diagnosed with Stage I-III invasive breast cancer between 2001-2011 who provided a blood sample. Individuals were excluded if they had prior cancer, except for non-melanoma skin cancer, and if they were missing information on lymph-node (LN) status.

#### Plasma fatty acid measurement

Measurement of plasma fatty acids was completed for a nested case-control sample within NHS2, including 794 breast cancer cases, and details have been reported^34^. Briefly, gas-liquid chromatography was used to measure erythrocyte fatty acid concentrations (in Dr. Hannia Campos’ laboratory at the Harvard T.H. Chan School of Public Health). A total of 39 fatty acids were analyzed and all measures were recorded as a percentage of total fatty acid. Measured monounsaturated fatty acids included: mysristoleic acid (14: 1n-5c), pentadecenoic acid (15:1n-5c), palmitoleic acid (16:1n-7c), oleic acid (18:1n-9c), octadecenoic (18:1n-7c), gondoic acid (20:1n-9c), and nervonic acid (24:1n-9c). Measured polyunsaturated fatty acids included n-3 [alpha-linolenic acid (ALA; 18:3n-3c), eicosapenaenoic acid (EPA; 20:5n-3c), docosapentaenoic acid (DPA; 22:5n-3c), and docosahexaenoic acid (DHA; 22:6n-3c)] and n-6 [linoleic acid (18:2n-6cc), gamma-linoleic acid (18:3n-6c), eicosadienoic acid (20:2n-6c), dihomo-gamma linolenic acid (20:3n-6c), arachidonic acid (20:4n-6c), docosadienoic acid (22:2n-6c), and aolrenic acid (22:4n-6c)] forms.

#### Metabolite profiling

Profiling of plasma metabolites was performed at the Broad Institute of MIT and Harvard (Cambridge, MA) via liquid chromatography tandem mass spectrometry (LC-MS/MS). Two platforms were used to identify metabolites – developed to measure polar metabolites (hydrophilic interaction liquid chromatography (HILIC-platform)), and lipids and free fatty acids (C8-positive platform), respectively – both of which have demonstrated good reproducibility and stability in prior publications^38^.

To allow for quality assessment, pooled reference samples were included (every 20 samples), and 64 quality controls were randomly distributed within batches. The relative abundance of each metabolite was determined by the integration of LC-MS peak areas, which are proportional to metabolite concentrations. For each metabolite, standardization was performed by taking the ratio of the sample to the nearest pooled reference and multiplying by the median of all reference values for the metabolite. For metabolites measured with multiple metabolomics platforms, the assay laboratory provided a list of the preferred measurement platforms. For metabolites measured multiple times with the same platform, the metabolite with the lowest CV was used for analysis and those with poor stability due to delay in processing^38^ (N=67) were excluded. Following this initial data cleaning, a total of 382 known metabolites were successfully measured and included in the study. Metabolites were annotated by class based on structural similarities.

Hydrophilic interaction liquid chromatography (HILIC) analyses of water soluble metabolites in the positive ionization mode were conducted using an LC-MS system com-prised of a Shimadzu Nexera X2 U-HPLC (Shimadzu Corp.; Marlborough, MA) coupled to a Q Exactive mass spectrometer (Thermo Fisher Scientific; Waltham, MA). Metabolites were extracted from plasma (10 µL) using 90 µL of acetonitrile/methanol/formic acid (74.9:24.9:0.2 v/v/v) containing stable isotope-labeled internal standards (valine-d8, Sig-ma-Aldrich; St. Louis, MO; and phenylalanine-d8, Cambridge Isotope Laboratories; And-over, MA). The samples were centrifuged (10 min, 9,000 x g, 4°C), and the supernatants were injected directly onto a 150 x 2 mm, 3 µm Atlantis HILIC column (Waters; Milford, MA). The column was eluted isocratically at a flow rate of 250 µL/min with 5% mobile phase A (10 mM ammonium formate and 0.1% formic acid in water) for 0.5 minute fol-lowed by a linear gradient to 40% mobile phase B (acetonitrile with 0.1% formic acid) over 10 minutes. MS analyses were carried out using electrospray ionization in the positive ion mode using full scan analysis over 70-800 m/z at 70,000 resolution and 3 Hz data acquisi-tion rate. Other MS settings were: sheath gas 40, sweep gas 2, spray voltage 3.5 kV, capil-lary temperature 350°C, S-lens RF 40, heater temperature 300°C, microscans 1, automatic gain control target 1e6, and maximum ion time 250 ms.

Plasma lipids were profiled using a Shimadzu Nexera X2 U-HPLC (Shimadzu Corp.; Marlborough, MA). Lipids were extracted from plasma (10 µL) using 190 µL of isopropa-nol containing 1,2-didodecanoyl-sn-glycero-3-phosphocholine (Avanti Polar Lipids; Ala-baster, AL). After centrifugation, supernatants were injected directly onto a 100 x 2.1 mm, 1.7 µm ACQUITY BEH C8 column (Waters; Milford, MA). The column was eluted isocrat-ically with 80% mobile phase A (95:5:0.1 vol/vol/vol 10mM ammonium ace-tate/methanol/formic acid) for 1 minute followed by a linear gradient to 80% mobile-phase B (99.9:0.1 vol/vol methanol/formic acid) over 2 minutes, a linear gradient to 100% mobile phase B over 7 minutes, then 3 minutes at 100% mobile-phase B. MS analyses were carried out using electrospray ionization in the positive ion mode using full scan analysis over 200–1100 m/z at 70,000 resolution and 3 Hz data acquisition rate. Other MS settings were: sheath gas 50, in source CID 5 eV, sweep gas 5, spray voltage 3 kV, capillary temperature 300°C, S-lens RF 60, heater temperature 300°C, microscans 1, automatic gain control target 1e6, and maximum ion time 100 ms. Lipid identities were denoted by total acyl carbon number and total double bond number.

Raw data from orbitrap mass spectrometers were processed using TraceFinder 3.3 software (Thermo Fisher Scientific; Waltham, MA) and Progenesis QI (Nonlinear Dynam-ics; Newcastle upon Tyne, UK). For analytical quality control, pooled plasma reference samples and mixtures of synthetic metabolite reference standards were analyzed at the beginning and end of sample queues to assure stable analytical performance, internal standard signals were evaluated in each sample to ensure consistent sample volume in-jections, and pooled plasma QC samples were inserted into the analytical queue at a fre-quency of 5% to evaluate analytical repeatability of each metabolite. Plasma samples were thawed on ice prior to aliquoting. As the aliquots for the LC-MS methods were prepared from each sample, a pooled plasma sample was created by placing an additional 10 μL aliquot from each sample into a 50 mL conical centrifuge tube. The pooled plasma sample was maintained on dry ice while samples were being aliquoted to promote rapid freezing and stored at −80 C in between sample batches until all additions were made. The pooled plasma was then thawed on ice, mixed by vortexing, and sub-aliquoted to create pooled plasma QC samples for each LC-MS method. For each method, metabolite identities were confirmed using mixtures of authentic reference standards (that were previously individ-ually identified in human plasma based on matching retention times, m/z, and MS/MS spectra) or reference samples (see Supplementary Table S10).

#### Covariates

Covariates included date (month), age (years), menopausal status (pre-vs. postmenopausal), postmenopausal hormone use (yes/no), and fasting status (≥10 hours since last meal vs. <10 hours since last meal) at time of blood draw, as well as established breast cancer risk factors. Established breast cancer risk factors were reported at the time of blood draw and/or at the most recent follow-up questionnaire preceding blood collection, and included: BMI at age 18y (kg/m^2^), weight change since age 18y (kg), age at first birth and parity (nulliparous, 1-2 children <25 y, 1-2 children 25+ y, 3+ children <25y, 3+ children 25+ y), age at menarche (years), breastfeeding history (yes/no), personal history of benign breast disease (yes/no), family history of breast cancer (yes/no), physical activity (MET-hours/week), alcohol intake (g/day), oral contraceptive use (yes/no), and race (White v. non-White). Missing covariates were imputed with either the most common value [age at menarche (13 years, N=5)], or the mean value [weight change (12.3 kg, N=2); BMI at age 18 (20.9 kg/m^2^, N=2); physical activity (18.1 met-hours/week, N=1); alcohol consumption (3.6 g/day, N=11)]. Parous individuals missing breastfeeding ever/never status (N=247) were assumed to have not breastfed.

### Statistical Analysis

#### Plasma MUFA/PUFA associations

We calculated the odds ratios (OR) and 95% confidence intervals (CI) for the association of plasma MUFA/PUFA ratio with LN-positivity using logistic regression. Models adjusted for: (1) blood draw factors only, (2) blood draw factors + BMI at age 18y + weight change since age 18y, and (3) full list of covariates (provided above). Odds ratios were reported to account for a change in the 10^th^ to 90^th^ percentile of MUFA/PUFA ratio, which was equal to 0.137 in this cohort.

#### Metabolite associations

All metabolites were probit transformed prior to analyses. Out of a total of 382 measured known metabolites, 294 (77%) had no missing values. Among the remaining 88 metabolites, those with <10% missingness (N=55) were imputed with ½ the minimum metabolite value and were used in primary analyses (N=349). Imputation was not performed on metabolites with ≥10% missing (N=33), and these metabolites were analyzed separately.

Among breast cancer cases, we calculated the odds ratios (OR) and 95% confidence intervals (CI) for the association of individual metabolite levels with lymph node positivity using logistic regression. Reported ORs were representative of a 2.5 standard deviation (SD) increase in metabolites, equivalent to the comparison of the 10^th^ percentile to the 90^th^ percentile of metabolite level under the assumption of a normal distribution. We used three different regression models for this assessment: (1) adjusting for blood draw factors only, (2) adding adjustment for BMI at age 18y and weight change since age 18y, and (3) including all covariates listed above. We additionally conducted a sensitivity analysis adding tumor subtype as an adjustment factor. In all analyses of individual metabolites, multiple correlated hypotheses were accounted for with adjustment by the number of effective tests, which was calculated by performing a principal components analysis of all metabolites among cases and calculating the number of principal components that explained 99.5% of the total variance^39^.

Average plasmalogen levels were calculated for each individual by taking the sum of probit-transformed PC and PE plasmalogens and dividing by the total number (N=25). We conducted a logistic regression analysis using average plasmalogen levels as the exposure of interest, with ORs representing a contrast between the 10^th^ and the 90^th^ percentile of the average value. The same models (1,2,3) described above were run. P-values were calculated directly within logistic regression models without need for multiple adjustment correction.

Metabolite set enrichment analysis (MSEA) was performed to test the combined influence of metabolites within the same structural subclass on LN positivity. Classes were defined by our collaborators at the Broad Institute of MIT and Harvard (Cambridge, MA) based on structural similarities. Triacylglycerols (TAGs) were further divided as TAGs with ≥3 vs. TAGs with <3 double bonds. MSEA has been described in detail elsewhere; briefly, after identifying metabolite subgroups, individual effect estimates (from logistic regressions) are combined and a summary Enrichment Score (ES), representing how much a metabolite group is enriched compared to other groups is calculated. A positive score indicates a positive enrichment of the metabolite set in LN-positive breast cancer cases. The Normalized Enrichment Score (NES) is further adjusted for group size, measured by the number of metabolites within the group^40^. P-values were adjusted using the False Discovery Rate (FDR) to account for multiple comparisons^41^. Datasets for analysis were created in SAS version 9 (SAS Institute Inc., Cary, NC, USA). All statistical analyses were conducted using the R programming language, version 4.0.3.

#### In vivo mouse dietary studies

For the murine metastasis experiments, we used a clinically relevant immunocompetent model of breast cancer metastasis, 10,000 4T1 cells injected into Balb/c mice. The 4T1 line spontaneously metastasizes after orthotopic injection into the mammary fat pad (LN-the “primary tumor” site, from which cancer cells spontaneously metastasize to lymph nodes) or after direct intranodal injection into the popliteal lymph node (LN+).

6-week-old female BALB/c immunodeficient mice were assigned into 4 dietary intervention groups. Mice were maintained for 7-8 weeks on isocaloric control group diet with varying MUFA/PUFA ratios by adjusting the proportions of specific fatty acids in each diet, linoleic acid enriched diet fat from safflower oil (Q1, MUFA/PUFA= 0.1), soybean oil (Q2, MUFA/PUFA= 0.4), palmitic acid enriched diet fat from palm oil (Q3, MUFA/PUFA= 3.3) and oleic acid enriched diet fat from safflower oil enriched with OA (Q4, MUFA/PUFA= 5.5). In each of these diets, the macronutrient composition was 20% protein, 55% carbohydrate, and 25% fat (Research Diets Inc, New Brunswick, NJ). Mice were housed 5 per cage with a 12:12 h light-dark cycle at 22–23 °C with free access to water and measured amounts of food. Mice were monitored regularly to ensure that they had sufficient water and food. Body weight and food intake were recorded weekly for 7-8 weeks after diet initiation. All animal protocols were approved by the Institutional Animal Care and Use Committee of Harvard University (IS00003460).

The mice after 2 weeks of dietary intervention were injected with 10,000 4T1 cells in the mammary fat pad and lymph node, and tumor formation data measured two times per week. Across all experiments, the maximum allowed tumor diameter was set at 2.0 cm, and this threshold was not surpassed in any instance. At that point, all mice in the cohort were euthanized per approved protocol, tumors were weighed and measured. Plasma, lymph fluid, tumor and lungs were collected, and cancer cells were isolated for furthered analysis and metastatic disease burden were evaluated by bioluminescence imaging of visceral organs. Blood was collected by cardiac puncture at week 7 into gel plasma separator microtainer tubes (BD microtainer capillary blood collector, Fisher Scientific) and centrifuged for plasma separation and stored at −80 °C for further analyses. Lymph fluids were extracted by collecting inguinal, axillary, and sciatic lymph nodes in a sterile plate on dry ice. The lymph nodes were then minced with a blade. Next, 100 µL of saline was added, and the mixture was pipetted up and down to facilitate the release of the fluid from the tissue. The tissue remnants were removed, the collected fluid was snap frozen for lymph fluid lipidomic analysis.

#### Cell lines

4T1 (ATCC® CRL2539™) murine breast cancer cells used in this study were purchased from the American Type Culture Collection ((ATCC), Manassas, VA, USA). Samples of 4T1 murine breast cancer cells were cultured in complete RPMI1640 medium containing 10% FBS, 1% penicillin (P) and 1% streptomycin (S). Cell lines were maintained in an incubator with 5% CO2 at 37 °C.

#### Intranodal cell injections

Murine lymphatics were first traced by injecting 2% Evans Blue Dye (product E2129, Sigma-Aldrich) into the footpad 5 minutes prior to the intranodal injections. Following the injection of Evans Blue Dye, the mice were anesthetized using isoflurane, and a small (5–10 mm) incision was made in the popliteal lymph node region. The lymph node, identified by the Evans Blue staining, was immobilized with forceps, 10,000 4T1 cells suspended in PBS were injected into the popliteal lymph node in a volume of 10μL using a Hamilton syringe. Successful injection into the lymph node was confirmed by visible swelling. The incision was then closed using surgical glue (product 1469SB, 3M VetBond Tissue Adhesive), and the mice were carefully monitored for any signs of pain or distress.

#### Orthotopic Mammary fat pad injections

Mice were anesthetized using isoflurane 10,000 4T1 cells suspended in 100 uL PBS per mouse were injected by a 27 gauge PrecisionGlide Needle (BD) guided ear the nipple. Carefully advance the needle tip into the subcutaneous mammary fat pad, positioning it just below the nipple confirmed by visible swelling of the skin, and the mice were carefully monitored for any signs of pain or distress.

#### Tumor digestion for flow cytometry

At the experimental endpoint, tumors from the mammary fat pad or from the lymph nodes were carefully dissected and weighted. Next, tumors were mechanically dislocated with a razor blade, one chunk was enzymatically digested and placed into 1.5mL of RPMI+10% FBS+1% Pen/Strep. Tumor were processed with warm digestion media containing collagenase I, hyaluronidase, DNase I and RPMI 1% FBS 1% PS and tissue was filtered through a 70µM strainer. Cells were then transferred to the FACS tubes stained with C11-BODIPY for flow cytometry by FACS Fortessa (BD Biosciences). The data was analyzed by FlowJo 10.8.1.

#### Analysis of lipid peroxidation by BODIPY-C11 staining

For lipid ROS detection using BODIPY-C11 staining, cells were suspended in 1 mL PBS supplemented with 5 μM BODIPY 665/676 (product B3832, Thermo Fisher Scientific) and incubated for 15 minutes at 37°C. Subsequently, cells were washed, resuspended in fresh PBS with 0.5 µg/ml DAPI, and immediately analyzed by flow cytometry. The median fluorescence intensity shifts from the non-oxidized (665 nm) to the oxidized (676 nm) channel was assessed to quantify BODIPY staining.

#### H&E analysis and microscopy

Lung tissues were fixed in zinc-formalin fixative (Z-Fix) for 48 hours and then transferred to 70% ethanol until processed for histological analysis. Following fixation, tissues were subjected to a standard dehydration protocol and subsequently embedded in paraffin. Sections approximately 5 µm thick were cut using a microtome and mounted on silane-coated glass slides. These slides were then incubated in a thermostat at 65°C for 1 hour to ensure proper adhesion of tissue sections to the slides. To prepare the sections for hematoxylin-eosin (H&E) staining, the slides were first dewaxed using xylene and rehydrated through a graded series of ethanol solutions, followed by a final rinse in distilled water. The H&E staining was performed by the Rodent Histology Department at Harvard Medical School, adhering to their established staining protocols to ensure consistent and high-quality results. Microscopic images of the stained sections were captured using a Nikon microscope equipped with a 10X objective lens (Nikon Eclipse Ni-e, 931245, Tokyo, Japan), Images were taken at random fields to ensure unbiased sampling by NIS-element software (AR5.42.03). Metastasis regions in both lungs were identified and quantified by counting the number of metastatic foci present in each section.

#### Metabolite extraction and measurements by LC/MS

The extraction buffer (acetonitrile: methanol: water, 4: 4: 2, v: v: v) was added to the lymph fluid or plasma at 1:20 ratio. The sample was homogenized by vortexing for 10 s. Then, the mixture was centrifuged (17,000 g, 10 min, 4 °C), and the supernatant was transferred to an LC-MS vial and analyzed on the LC-MS. To facilitate quality assessment, a pooled quality control (QC) sample was generated by combining 20 µL from each experimental sample and was injected every five samples during the analytical run.

Metabolite extracts were analyzed using a quadrupole-orbitrap mass spectrometer coupled with hydrophilic interaction chromatography (HILIC). Chromatographic separation was achieved on an XBridge BEH Amide XP Column (2.5 µm, 2.1 mm × 150 mm) with a guard column (2.5 µm, 2.1 mm × 5 mm) (Waters, Milford, MA). Chromatography was conducted on a Vanquish™ UHPLC system (Thermo Scientific, Waltham, MA), using mobile phase A (water:acetonitrile 95:5) and mobile phase B (water:acetonitrile 20:80), both containing 10 mM ammonium acetate and 10 mM ammonium hydroxide. The linear elution gradient was: 0 ∼ 3 min, 100% B; 3.2 ∼ 6.2 min, 90% B; 6.5. ∼ 10.5 min, 80% B; 10.7 ∼ 13.5 min, 70% B; 13.7 ∼ 16 min, 45% B; and 16.5 ∼ 22 min, 100% B, with a flow rate of 0.3 mL/ min. The autosampler was at 4°C. The column temperature was 30 °C. The injection volume was 5 µL. Needle wash was applied between samples using acetonitrile: methanol: water at 4: 4: 2 (v: v: v). For mass spectrometry, an Orbitrap Exploris™ 480 Mass Spectrometer (Thermo Scientific, Waltham, MA, USA) was used. For MS1 acquisition, the MS scanned from 70 to 1000 m/z with switching polarity at a resolution of 120,000 for all experimental samples. The relevant parameters were sheath gas, 40; auxiliary gas, 10; sweep gas, 2; spray voltage, 3.5 kV; capillary temperature, 300 °C; S-lens, 45. The resolution was set at 120,000 (at m/z 200). Maximum injection time (max IT) was set at 500 ms, and automatic gain control (AGC) was set at 3 × 10^6^.

MS1 raw data files were converted into mzxML using msconvert and imported to EI-Maven (Elucidata, Cambridge, MA) for targeted metabolomics. Metabolites were identified based on accurate mass and retention time with an in-house library.

#### Lipidomic extraction and measurements by LC/MS

The SPLASH™ LIPIDOMIX™ Mass Spec Standard (Avanti Polar Lipids, Birmingham, AL), containing 14 deuterated exogenous lipid standards in methanol, was used as an internal standard and spiked to the extraction buffer at a 1:20 ratio before use. Lipid extraction was performed as previously described^42,42^. The extraction buffer (butanol/methanol, 1:1, with 5 mM ammonium formate) containing the internal standard was added to lymph fluid or plasma at a 1:10 ratio. Samples were homogenized by vortexing for 1 min, followed by shaking for 30 min at 4 °C. After centrifugation (17,000 g, 10 min, 4 °C), the supernatant was transferred to glass LC-MS vials for analysis. For quality assessment, a pooled quality control (QC) sample was prepared by combining 20 µL from each experimental sample and injected every five to six samples during the analytical run.

Lipid extracts were analyzed using a Dionex Ultimate 3000 RSLC system coupled with a QExactive™ mass spectrometer (Thermo Scientific, Waltham, MA, USA). Chromatographic separation was achieved on an ACQUITY UPLC CSH C18 column (130Å, 1.7 µm, 2.1 mm × 100 mm) with an ACQUITY UPLC CSH C18 VanGuard pre-column (130Å, 1.7 µm, 2.1 mm × 5 mm) (Waters, Milford, MA) with column temperature at 50 °C. For the gradient, mobile phase A consisted of an acetonitrile-water mixture (6:4), and mobile phase B was a 2-propanol-acetonitrile mixture (9:1), both phases contained 10 mM ammonium formate and 0.1% formic acid. The linear elution gradient was: 0-3 min, 20% B; 3-7 min, 20-55% B; 7-15 min, 55-65% B; 15-21 min, 65-70% B; 21-24 min, 70-100% B; and 24-26 min, 100% B, 26-28 min, 100-20% B, 28-30 min, 20% B, with a flow rate of 0.35 mL/ min. The autosampler was at 4°C. The injection volume was 5 µL. Needle wash was applied between samples using a mixture of dichloromethane-isopropanol-acetonitrile (1:1:1).

ESI-MS analysis was performed in positive and negative ionization polarities using a combined full mass scan and data-dependent MS/MS (Top 10) (Full MS/dd-MS2) approach. The experimental conditions for full scanning were as follows: resolving power, 70,000; automatic gain control (AGC) target, 1 × 10^6^; and maximum injection time (IT), 100 ms. The scan range of the instrument was set to m/z 100-1200 in both positive and negative ion modes. The experimental conditions for the data-dependent product ion scanning were as follows: resolving power, 17,500; AGC target, 5 × 10^4^; and maximum IT, 50 ms. The isolation width and stepped normalized collision energy (NCE) were set to 1.0 m/z, and 10, 20, and 40 eV. The intensity threshold of precursor ions for dd-MS2 analysis and the dynamic exclusion were set to 1.6 × 105 and 10 s. The ionization conditions in the positive mode were as follows: sheath gas flow rate, 50 arb; auxiliary (AUX) gas flow rate, 15 arb; sweep gas flow rate, 1 arb; ion spray voltage, 3.5 kV; AUX gas heater temperature, 325 °C; capillary temperature, 350 °C; and S-lens RF level, 55. The ionization conditions in the negative mode were as follows: sheath gas flow rate, 45 arb; auxiliary (AUX) gas flow rate, 10 arb; sweep gas flow rate, 1 arb; ion spray voltage, 2.5 kV; AUX gas heater temperature, 320 °C; capillary temperature, 320 °C; and S-lens RF level, 55.

Thermo Scientific™ LipidSearch™ software version 5.0 was used for lipid identification and quantitation. First, the product search mode was used during which lipids were identified based on the exact mass of the precursor ions and the mass spectra resulting from product ion scanning. The precursor and product tolerances were set to 10 and 10 ppm mass windows. The absolute intensity threshold of precursor ions and the relative intensity threshold of product ions were set to 30000 and 1%. Next, the search results from the individual positive or negative ion files from each sample were aligned within a retention time window (±0.25 min) and then all the data were merged for each annotated lipid with a retention time correction tolerance of 0.5 min. The annotated lipids were then filtered to reduce false positives by only including the lipids with a total grade of A or B.

## Acknowledgements

This work was supported by the Ludwig Cancer Center at Harvard (J.M.U.), NIH NCI R01CA282202 (J.M.U., O.A.Z subcontract). Research was supported by NIH NCI U01 CA176726 (A.H.E.). The content is solely the responsibility of the authors and does not necessarily represent the official views of the National Institutes of Health. The authors would like to acknowledge the contribution to this study from central cancer registries supported through the Centers for Disease Control and Prevention’s National Program of Cancer Registries (NPCR) and/or the National Cancer Institute’s Surveillance, Epidemiology, and End Results (SEER) Program. Central registries may also be supported by state agencies, universities, and cancer centers. Participating central cancer registries include the following: Alabama, Alaska, Arizona, Arkansas, California, Delaware, Colorado, Connecticut, Florida, Georgia, Hawaii, Idaho, Indiana, Iowa, Kentucky, Louisiana, Maine, Maryland, Massachusetts, Michigan, Mississippi, Montana, Nebraska, Nevada, New Hampshire, New Jersey, New Mexico, New York, North Carolina, North Dakota, Ohio, Oklahoma, Oregon, Pennsylvania, Puerto Rico, Rhode Island, Seattle SEER Registry, South Carolina, Tennessee, Texas, Utah, Virginia, West Virginia, Wyoming.

## Author contributions

M.Y. designed the in vivo study, performed the experiments, and analyzed the data. K.D.B. conducted the NHS data analysis and data interpretation and wrote the NHS manuscript section. A.C. assisted with in vivo endpoint experiments and lymph node collection. Y.L. conducted mass spectrometry runs. J.M.L. designed initial lipidomic data analysis. M.S. provided input on manuscript writing and assisted with the in vivo study. J.T., K.J.S., M.F., M.P., and C.S.F. assisted with the in vivo study. S.T.H. supervised the mass spectrometry core. W.H., W.C.W., and A.H.E. provided input on manuscript writing. O.A.Z. and J.M.U. supervised the study.

All authors reviewed and approved the final version of the manuscript.

## References

1. Practice Guidelines in Oncology: Breast Cancer. 2021: National Comprehensive Cancer Network (NCCN). Accessed at https://www.nccn.org/guidelines/ on April 7, 2024.

2. Boire A, Burke K, Cox TR, Guise T, Jamal-Hanjani M, Janowitz T, Kaplan R, Lee R, Swanton C, Vander Heiden MG, Sahai E. Why do patients with cancer die? Nat Rev Cancer. 2024 Aug;24(8):578–589. doi: 10.1038/s41568-024-00708-4. Epub 2024 Jun 19. PMID: 38898221; PMCID: PMC7616303.

3. Mani K, Deng D, Lin C, Wang M, Hsu ML, Zaorsky NG. Causes of death among people living with metastatic cancer. Nat Commun. 2024 Feb 19;15(1):1519. doi: 10.1038/s41467-024-45307-x. PMID: 38374318; PMCID: PMC10876661.

4. Brown M, Assen FP, Leithner A, Abe J, Schachner H, Asfour G, Bago-Horvath Z, Stein JV, Uhrin P, Sixt M, Kerjaschki D. Lymph node blood vessels provide exit routes for metastatic tumor cell dissemination in mice. Science. 2018;359(6382):1408–11. doi: 10.1126/science.aal3662. PubMed PMID: 29567714.

5. Pereira ER, Kedrin D, Seano G, Gautier O, Meijer EFJ, Jones D, Chin SM, Kitahara S, Bouta EM, Chang J, Beech E, Jeong HS, Carroll MC, Taghian AG, Padera TP. Lymph node metastases can invade local blood vessels, exit the node, and colonize distant organs in mice. Science. 2018;359(6382):1403–7. Epub 20180322. doi: 10.1126/science.aal3622. PubMed PMID: 29567713; PMCID: PMC6002772.

6. Sleeman J, Schmid A, Thiele W. Tumor lymphatics. Semin Cancer Biol. 2009;19(5):285-97. Epub 20090529. doi: 10.1016/j.semcancer.2009.05.005. PubMed PMID: 19482087.

7. Alitalo A, Detmar M. Interaction of tumor cells and lymphatic vessels in cancer progression. Oncogene. 2012;31(42):4499–508. Epub 20111219. doi: 10.1038/onc.2011.602. PubMed PMID: 22179834.

8. Jatoi I, Hilsenbeck SG, Clark GM, Osborne CK. Significance of axillary lymph node metastasis in primary breast cancer. J Clin Oncol. 1999 Aug;17(8):2334–40. doi: 10.1200/JCO.1999.17.8.2334. Erratum in: J Clin Oncol 1999 Oct;17(10):3365. PMID: 10561295.

9. Wilking N, Rutqvist LE, Carstensen J, Mattsson A, Skoog L. Prognostic significance of axillary nodal status in primary breast cancer in relation to the number of resected nodes. Stockholm Breast Cancer Study Group. Acta Oncol. 1992;31(1):29–35. doi: 10.3109/02841869209088261. PMID: 1586501.

10. Lee CK, Jeong SH, Jang C, Bae H, Kim YH, Park I, Kim SK, Koh GY. Tumor metastasis to lymph nodes requires YAP-dependent metabolic adaptation. Science. 2019;363(6427):644–9. Epub 20190207. doi:10.1126/science.aav0173. PubMed PMID: 30733421.

11. Martin-Perez M, Urdiroz-Urricelqui U, Bigas C, Benitah SA. The role of lipids in cancer progression and metastasis. Cell Metab. 2022 Nov 1;34(11):1675–1699. doi: 10.1016/j.cmet.2022.09.023. Epub 2022 Oct 18. PMID: 36261043.

12. Pascual G, Avgustinova A, Mejetta S, Martín M, Castellanos A, Attolini CS, Berenguer A, Prats N, Toll A, Hueto JA, Bescós C, Di Croce L, Benitah SA. Targeting metastasis-initiating cells through the fatty acid receptor CD36. Nature. 2017 Jan 5;541(7635):41–45. doi: 10.1038/nature20791. Epub 2016 Dec 7. PMID: 27974793.

13. Ubellacker JM, Tasdogan A, Ramesh V, Shen B, Mitchell EC, Martin-Sandoval MS, Gu Z, McCormick ML, Durham AB, Spitz DR, Zhao Z, Mathews TP, Morrison SJ. Lymph protects metastasizing melanoma cells from ferroptosis. Nature. 2020 Sep;585(7823):113–118. doi: 10.1038/s41586-020-2623-z. Epub 2020 Aug 19. PMID: 32814895; PMCID: PMC7484468.

14. Magtanong L, Ko PJ, To M, Cao JY, Forcina GC, Tarangelo A, Ward CC, Cho K, Patti GJ, Nomura DK, Olzmann JA, Dixon SJ. Exogenous Monounsaturated Fatty Acids Promote a Ferroptosis-Resistant Cell State. Cell Chem Biol. 2019 Mar 21;26(3):420–432.e9. doi: 10.1016/j.chembiol.2018.11.016. Epub 2019 Jan 24. PMID: 30686757; PMCID: PMC6430697.

15. Brantley KD, Zeleznik OA, Rosner B, Tamimi RM, Avila-Pacheco J, Clish CB, Eliassen AH. Plasma Metabolomics and Breast Cancer Risk over 20 Years of Follow-up among Postmenopausal Women in the Nurses’ Health Study. Cancer Epidemiol Biomarkers Prev. 2022;31(4):839–50. doi: 10.1158/1055-9965.EPI-21-1023. PubMed PMID: 35064065; PMCID: PMC8983458.

16. Zeleznik OA, Balasubramanian R, Ren Y, Tobias DK, Rosner BA, Peng C, Bever AM, Frueh L, Jeanfavre S, Avila-Pacheco J, Clish CB, Mora S, Hu FB, Eliassen AH. Branched-Chain Amino Acids and Risk of Breast Cancer. JNCI Cancer Spectr. 2021;5(5). Epub 20210702. PubMed PMID: 34585062; PMCID:PMC8460878.

17. Zeleznik OA, Balasubramanian R, Zhao Y, Frueh L, Jeanfavre S, Avila-Pacheco J, Clish CB, Tworoger SS, Eliassen AH. Circulating amino acids and amino acid-related metabolites and risk of breast cancer among predominantly premenopausal women. NPJ Breast Cancer. 2021;7(1):54. Epub 20210518. doi: 10.1038/s41523-021-00262-4. PubMed PMID: 34006878; PMCID: PMC8131633.

18. Sabatier M, Solanki A, Thangaswamy S, Lei PJ, Zhou H, O’Melia M, Menzel L, Mitri S, Ubellacker JM. Lymphatic collection and cell isolation from mouse models for multiomic profiling. Nat Protoc. 2025 Feb 10. doi: 10.1038/s41596-025-01156-6. PMID: 39779897.

19. Dixon SJ, Lemberg KM, Lamprecht MR, Skouta R, Zaitsev EM, Gleason CE, Patel DN, Bauer AJ, Cantley AM, Yang WS, Morrison B 3rd, Stockwell BR. Ferroptosis: an iron-dependent form of nonapoptotic cell death. Cell. 2012 May 25;149(5):1060–72. doi: 10.1016/j.cell.2012.03.042. PMID: 22632970; PMCID: PMC3367386.

20. Doll S, Proneth B, Tyurina YY, Panzilius E, Kobayashi S, Ingold I, Irmler M, Beckers J, Aichler M, Walch A, Prokisch H, Trümbach D, Mao G, Qu F, Bayir H, Füllekrug J, Scheel CH, Wurst W, Schick JA, Kagan VE, Angeli JP, Conrad M. ACSL4 dictates ferroptosis sensitivity by shaping cellular lipid composition. Nat Chem Biol. 2017 Jan;13(1):91–98. doi: 10.1038/nchembio.2239. Epub 2016 Nov 14. PMID: 27842070; PMCID: PMC5610546.

21. Kagan VE, Mao G, Qu F, Angeli JP, Doll S, Croix CS, Dar HH, Liu B, Tyurin VA, Ritov VB, Kapralov AA, Amoscato AA, Jiang J, Anthonymuthu T, Mohammadyani D, Yang Q, Proneth B, Klein-Seetharaman J, Watkins S, Bahar I, Greenberger J, Mallampalli RK, Stockwell BR, Tyurina YY, Conrad M, Bayır H. Oxidized arachidonic and adrenic PEs navigate cells to ferroptosis. Nat Chem Biol. 2017 Jan;13(1):81–90. doi: 10.1038/nchembio.2238. Epub 2016 Nov 14. PMID: 27842066; PMCID: PMC5506843.

22. de Oliveira ST, Bessani MP, Scandolara TB, Silva JC, Kawassaki ACB, Fagotti PAF, Maito VT, de Souza JA, Rech D, Panis C. Systemic lipid peroxidation profile from patients with breast cancer changes according to the lymph nodal metastasis status. Oncoscience. 2022 Feb 24;9:1–10. doi: 10.18632/oncoscience.550. PMID: 35233438; PMCID: PMC8876690.

23. Verma N, Vinik Y, Saroha A, Nair NU, Ruppin E, Mills G, Karn T, Dubey V, Khera L, Raj H, Maina F, Lev S. Synthetic lethal combination targeting BET uncovered intrinsic susceptibility of TNBC to ferroptosis. Sci Adv. 2020 Aug 21;6(34):eaba8968. doi: 10.1126/sciadv.aba8968. PMID: 32937365; PMCID: PMC7442484.

24. Nagan N, Zoeller RA. Plasmalogens: biosynthesis and functions. Prog Lipid Res. 2001 May;40(3):199-229. doi: 10.1016/s0163-7827(01)00003-0. PMID: 11275267.

25. Braverman NE, Moser AB. Functions of plasmalogen lipids in health and disease. Biochim Biophys Acta. 2012 Sep;1822(9):1442–52. doi: 10.1016/j.bbadis.2012.05.008. Epub 2012 May 22. PMID: 22627108.

26. Wu Y, Chen Z, Jia J, Chiba H, Hui SP. Quantitative and Comparative Investigation of Plasmalogen Species in Daily Foodstuffs. Foods. 2021 Jan 8;10(1):124. doi: 10.3390/foods10010124. PMID: 33435634; PMCID: PMC7827193.

27. Han X, Holtzman DM, McKeel DW Jr. Plasmalogen deficiency in early Alzheimer’s disease subjects and in animal models: molecular characterization using electrospray ionization mass spectrometry. J Neurochem. 2001 May;77(4):1168–80. doi: 10.1046/j.1471-4159.2001.00332.x. PMID: 11359882.

28. Fabelo N, Martín V, Santpere G, Marín R, Torrent L, Ferrer I, Díaz M. Severe alterations in lipid composition of frontal cortex lipid rafts from Parkinson’s disease and incidental Parkinson’s disease. Mol Med. 2011 Sep-Oct;17(9-10):1107-18. doi: 10.2119/molmed.2011.00119. Epub 2011 Jun 22. PMID: 21717034; PMCID: PMC3188884.

29. Zou Y, Henry WS, Ricq EL, Graham ET, Phadnis VV, Maretich P, Paradkar S, Boehnke N, Deik AA, Reinhardt F, Eaton JK, Ferguson B, Wang W, Fairman J, Keys HR, Dančík V, Clish CB, Clemons PA, Hammond PT, Boyer LA, Weinberg RA, Schreiber SL. Plasticity of ether lipids promotes ferroptosis susceptibility and evasion. Nature. 2020 Sep;585(7826):603–608. doi: 10.1038/s41586-020-2732-8. Epub 2020 Sep 16. PMID: 32939090; PMCID: PMC8051864.

30. Cui W, Liu D, Gu W, Chu B. Peroxisome-driven ether-linked phospholipids biosynthesis is essential for ferroptosis. Cell Death Differ. 2021 Aug;28(8):2536–2551. doi: 10.1038/s41418-021-00769-0. Epub 2021 Mar 17. PMID: 33731874; PMCID: PMC8329287.

31. Perez MA, Clostio AJ, Houston IR, Ruiz J, Magtanong L, Dixon SJ, Watts JL. Ether lipid deficiency disrupts lipid homeostasis leading to ferroptosis sensitivity. PLoS Genet. 2022 Sep 30;18(9):e1010436. doi: 10.1371/journal.pgen.1010436. PMID: 36178986; PMCID: PMC9555615.

32. Colditz G. A. & Hankinson S. E. The Nurses’ Health Study: lifestyle and health among women. Nat Rev Cancer 5, 388–396 (2005).

33. Tworoger SS, Sluss P, Hankinson SE (2006) Association between plasma prolactin concentrations and risk of breast cancer among predominately premenopausal women. Can Res 66(4):2476–2482

34. Hirko KA, Chai B, Spiegelman D, et al. Erythrocyte membrane fatty acids and breast cancer risk: a prospective analysis in the nurses’ health study II. Int J Cancer. 2018;142(6):1116–1129. doi:10.1002/ijc.31133

35. Mascanfroni ID, Takenaka MC, Yeste A, Patel B, Wu Y, Kenison JE, et al. Metabolic control of type 1 regulatory T cell differentiation by AHR and HIF1-α. Nat Med. 2015;21(6):638–646. doi: 10.1038/nm.3868

36. O’Sullivan JF, Morningstar JE, Yang Q, Zheng B, Gao Y, Jeanfavre S, et al. Dimethylguanidino valeric acid is a marker of liver fat and predicts diabetes. J Clin Invest. 2017;127(12):4394–4402. doi: 10.1172/JCI95995

37. Paynter NP, Balasubramanian R, Giulianini, Wang DD, Tinker LF, Gopal S, et al., Metabolic predictors of incident coronary heart disease in women. Circulation, 2018. 137(8): p. 841–853.

38. Townsend MK, Clish CB, Kraft P, Wu C, Souza AL, Deik AA, et al. Reproducibility of metabolomic profiles among men and women in 2 large cohort studies. Clin Chem. 2013;59:1657–67. doi: 10.1373/clinchem.2012.199133.39.

39. Gao X., Starmer J., Martin E.R. A multiple testing correction method for genetic association studies using correlated single nucleotide polymorphisms. Genet Epidemiol. 2008;32:361–369. doi: 10.1002/gepi.20310.

40. Subramanian A, Tamayo P, Mootha VK, Mukherjee S, Ebert BL, Gillette MA, et al. Gene set enrichment analysis: a knowledge-based approach for interpreting genome-wide expression profiles. Proceedings of the National Academy of Sciences. 2005;102(43):15545–50.

41. Benjamini Y., Drai D., Elmer G., et al. Controlling the false discovery rate in behavior genetics research. Behav Brain Res. 2001;125:279–284. doi: 10.1016/s0166-4328(01)00297-2.

42. Alshehry ZH, Barlow CK, Weir JM, Zhou Y, McConville MJ, Meikle PJ. An Efficient Single Phase Method for the Extraction of Plasma Lipids. Metabolites. 2015 Jun 17;5(2):389–403. doi: 10.3390/metabo5020389. PMID: 26090945; PMCID: PMC4495379.

43. Wong MWK, Braidy N, Pickford R, Sachdev PS, Poljak A. Comparison of Single Phase and Biphasic Extraction Protocols for Lipidomic Studies Using Human Plasma. Front Neurol. 2019 Aug 21;10:879. doi: 10.3389/fneur.2019.00879. PMID: 31496985; PMCID: PMC6712511.

